# Role of HP1β during spermatogenesis and DNA replication

**DOI:** 10.1101/2019.12.22.886424

**Authors:** Vijaya Charaka, Anjana Tiwari, Raj K Pandita, Clayton R Hunt, Tej K. Pandita

**Author notes:** **Authors e-mail address:** Vijay K. Charaka, Anjana Tiwari, Raj K. Pandita, Clayton R Hunt, Tej K. Pandita. **Correspondence:** Tej K. Pandita.

## Abstract

Maintaining genomic stability in a continually dividing cell population requires accurate DNA repair, especially in male germ cells. Repair and replication protein access to DNA, however, is complicated by chromatin compaction. The HP1β chromatin protein, encoded by *Cbx1*, is associated with chromatin condensation but its role in meiosis is not clear. To investigate the role of *Cbx*1 in male germ cells, we generated testis specific *Cbx*1 deficient transgenic mice by crossing *Cbx*1^flox/flox^ (*Cbx*1^f/f^) mice with Stra8 *Cre*^+/−^ mice. Loss of *Cbx*1 in testes adversely affected sperm maturation and *Cbx*1 deletion increased seminiferous tubule degeneration and basal level DNA damage., We observed that *Cbx*1^−/−^ MEF cells displayed reduced resolution of stalled DNA replication forks as well as decreased fork restart, indicating defective DNA synthesis. Taken together, these results suggest that loss of *Cbx*1 in growing cells leads to DNA replication defects and associated DNA damage that impact cell survival.

## Introduction

Meiosis generates haploid daughter cells and normal meiotic progression depends upon the fidelity of homologous chromosomes undergoing pairing, synapsis, recombination and proper segregation. Specifically DNA synthesis with proper fidelity is essential for repair of programmed DNA double strand breaks (DSBs) that are essential for chromosome dynamics in meiotic prophase I. Meiotic DSBs are repaired through two pathways, a synthesis-dependent strand annealing (SDSA) pathway to form non-crossover products and the classical pathway through double Holiday Junction (dHJ) formation to create both crossover and non-crossover products (Fishel, 2015; Lu and Yu, 2015). The DSBs that persist till pachynema are more likely to be repaired through the classical pathway to generate crossovers (Allers and Lichten, 2001).

Gametogenesis results in a large number of base mismatches, single strand and double stranded DNA breaks being generated which are repaired through BER, MMR, HR and NHEJ pathways (Garcia-Rodriguez et al., 2018). Unlike mitotic cells, spermatogonia cells transition through phases of mitotic and meiotic cell divisions. In the mitotic phase, spermatogonia cells undergo series of DNA duplications resulting in spermatogonia B cells. Duplicated spermatogonia B cells then undergo a multi-step meiotic cell division to produce mature, haploid spermatocytes. In dividing cells HR predominantly occurs in mitotic S phase cells to maintain DNA fidelity During meiosis phase 1 the many double strand breaks generated during crossover are also repaired by HR (Cohen and Pollard, 2001; Roeder, 1997). Loss of HR repair related proteins, therefore, (such as 53BP1, RIF1, BRCA1, ATM, MOF) affects spermatogenesis (Bartkova et al., 2001; Jiang et al., 2018; Marcet-Ortega et al., 2017; Pandita et al., 1999; Schwab et al., 2013; Sun et al., 2016). *Cbx*1^−/−^ mice display prenatal death due to severe alveoli lung damage and abnormal neuromuscular and cerebral cortex tissue development (Aucott et al., 2008). Previously we reported that *Cbx*1^−/−^ MEFs have increased genomic instability (Aucott et al., 2008). To circumvent this *Cbx1*^−/−^ perinatal lethality, we crossed *Cbx*1^f/f^ mice with *Str8 Cre*^+/−^ mice to generate mice with testis germ cell *Cbx1* deletion. Testis deleted for *Cbx*1 were viable, but had abnormalities in spermatogenesis. *Cbx*1 loss in testis arrested spermatogonia cells at spermatogonia B stage and also increased basal level DNA damage. Damage during the early stages of spermatogenesis correlated with decreased sperm production, which caused a subfertility phenotype.

The increased DNA damage in replicating spermatogonia B cells due to loss of *Cbx*1 is possibly the result of defective DNA synthesis as we observed that *Cbx*1 MEF knock out or knockdown in HeLa by specific siRNA results in the defective replication fork progression and increased stalled fork repair.

## Results

### *Cbx*1 conditional knockout mice

To elucidate the *in vivo* function of HP1β (*Cbx*1) in rapidly dividing cells we examined testis germ cells by deleting the LacZ /Neomycin gene region in *Cbx1*^tm1a^ mice by crossing with FLPeR (flipper) mice (Farley et al., 2000). Homozygous *Cbx*1^f/f^ mice were generated by backcrossing the *Cbx*1^f/+^/*Frt*^+/−^ mice (Fig.1a) and tissue specific *Cbx*1 knock out in testis then generated by crossing *Cbx*1^f/f^ mice with *Stra8 Cre*^+/−^ mice using the strategy described previously (Gupta et al., 2013; Kumar et al., 2011). Cre expression deleted the second exon and shifted *Cbx*1 gene reading frame, which ultimately resulted in loss of functional HP1β protein in testis (Fig.1a).

**Figure 1.**
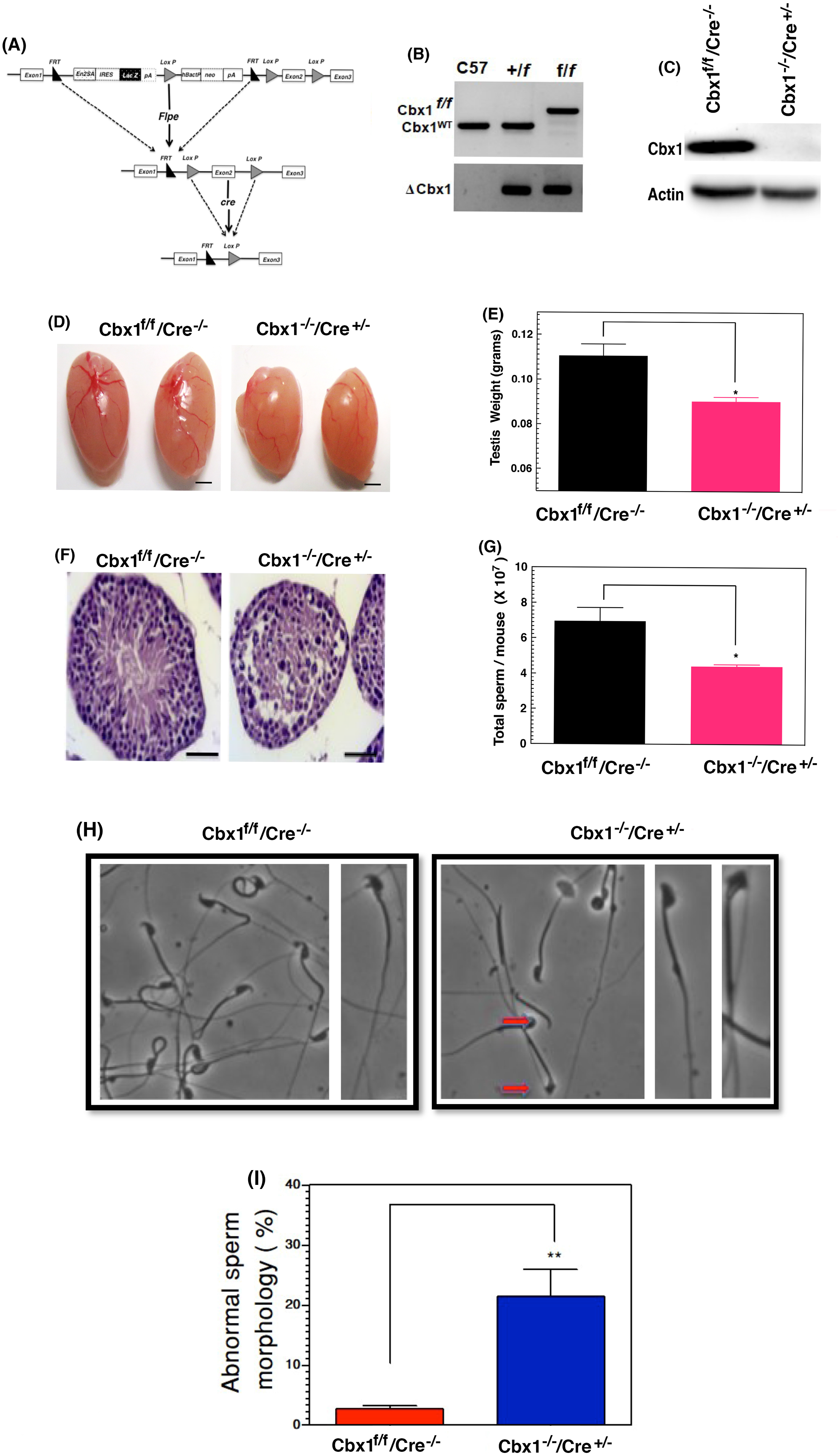
Phenotypes of *Cbx1* testis specific conditional KO mice. **(a)** Schematic map of the *Cbx*1 genomic locus surrounding the targeted exon. **(b)** PCR genotype analysis of the *Cbx1* WT and *Cbx1* testis specific conditional KO mice. **(c)** Western blot of protein extracts from *Cbx1* WT and *Cbx1* testis specific conditional KO mice with HP1β antibody. **(d)** Photographs of 6 month old *Cbx1* WT and *Cbx1* testis specific conditional KO mice testes are shown. **(e)** Measured weights of testis from 6 month *Cbx1* WT and *Cbx1* testis specific conditional KO mice (n=15). **(f)** H&E stained sections from 6 month old *Cbx1* testis specific conditional KO mice showing tubular vacuolation. **(g)** Sperm counts in epididymis from 6 months old *Cbx1* WT and *Cbx1* testis specific conditional KO mice, (n=15). **(h)** Sperm with typical hook-shaped head in contrast to round or irregular banana heads in *Cbx1* testis specific conditional KO mice. **(I)** Average mean of 15 male mice’s abnormal morphology sperm quantitated in *Cbx1* WT and *Cbx1* testis specific conditional KO mice. Data are analyzed and presented as mean ± SEM. * P<0.05, Student’s t-test.

Deletion of testis specific *Cbx*1 (*Cbx*1^−/−^/*Cre*^+/−^) in mice was confirmed by PCR (Fig.1b) and western blot analysis (Fig. 1c). All *Cbx*1^f/f^/*Cre*^−/−^ (*Cbx1* WT), *Cbx*1^f/+^/*Cre*^+/−^ (*Cbx*1 het) and conditional testis specific *Cbx*1^−/−^/*Cre*^+/−^ (*Cbx*1 testis specific conditional KO) mice survived. *Cbx1* WT and *Cbx1* het male mice displayed similar fertility levels, whereas in *Cbx1* testis specific conditional KO mice the litter size was half of *Cbx1* WT mice. *Cbx1* testis specific conditional KO mice had an average litter size of 3.07 and a 1/1.7 male/female ratio (Table 1) while wild-type mice had 7.87 pups per litter and a 1/1.28 male/female ratio. Thee results suggest testis specific loss of *Cbx1* caused a sub-fertility effect.

### HP1β is required for spermatogenesis and sperm maturation

The testis size of *Cbx1* WT and *Cbx*1 testis specific conditional KO mice was compared and the found to have testes about 20% smaller in size than testis of *Cbx1* WT mice (Fig.1d,e) and weighed less than *Cbx1* WT testis (Fig.1e). Since loss of DNA repair proteins has been linked with germ cell loss and seminiferous tubules degeneracy (Gunes et al., 2015; Xu et al., 1996), we examined whether *Cbx*1 loss in testis is causing similar defects. Tubular vacuolation and partial degeneration of spermatocytes was increased in *Cbx1* testis specific conditional KO mouse testes (Fig. 1f) and probably contributes to the observed reduced fertility.

Degeneration in seminiferous tubules is known to affect various stages of spermatogenesis and mature sperm production (Vyas et al., 2013). We harvested epididymis sperm from *Cbx1* WT and *Cbx1* testis specific conditional KO mice and found that *Cbx*1 loss had reduced sperm counts by 50% as compared to control mice (Fig. 1g). Moreover, the *Cbx*1 testis specific conditional KO mice had a 5-fold increase in abnormal sperms. *Cbx1* testis specific KO mice showed increase in sperms lacking hook and folded banana shaped head morphology (Fig. 1h,I). These results suggest that HP1β plays a role till pachytene stage of meiosis, as we did not observe HP1β in either wild-type or *Cbx1* testis specific conditional KO mouse testes post pachytene stage by immunofluorescence analysis (Fig. 2a). To investigate HP1β function in spermatogenesis, spermatocytes were prepared from *Cbx1* WT and *Cbx1* conditional KO testes and stained for synaptonemal complex protein 3 (SYCP3) revealed the loss of HP1β increased spermatocyte arrest at the spermatogonia stage (Fig. 2b,c).

**Figure 2.**
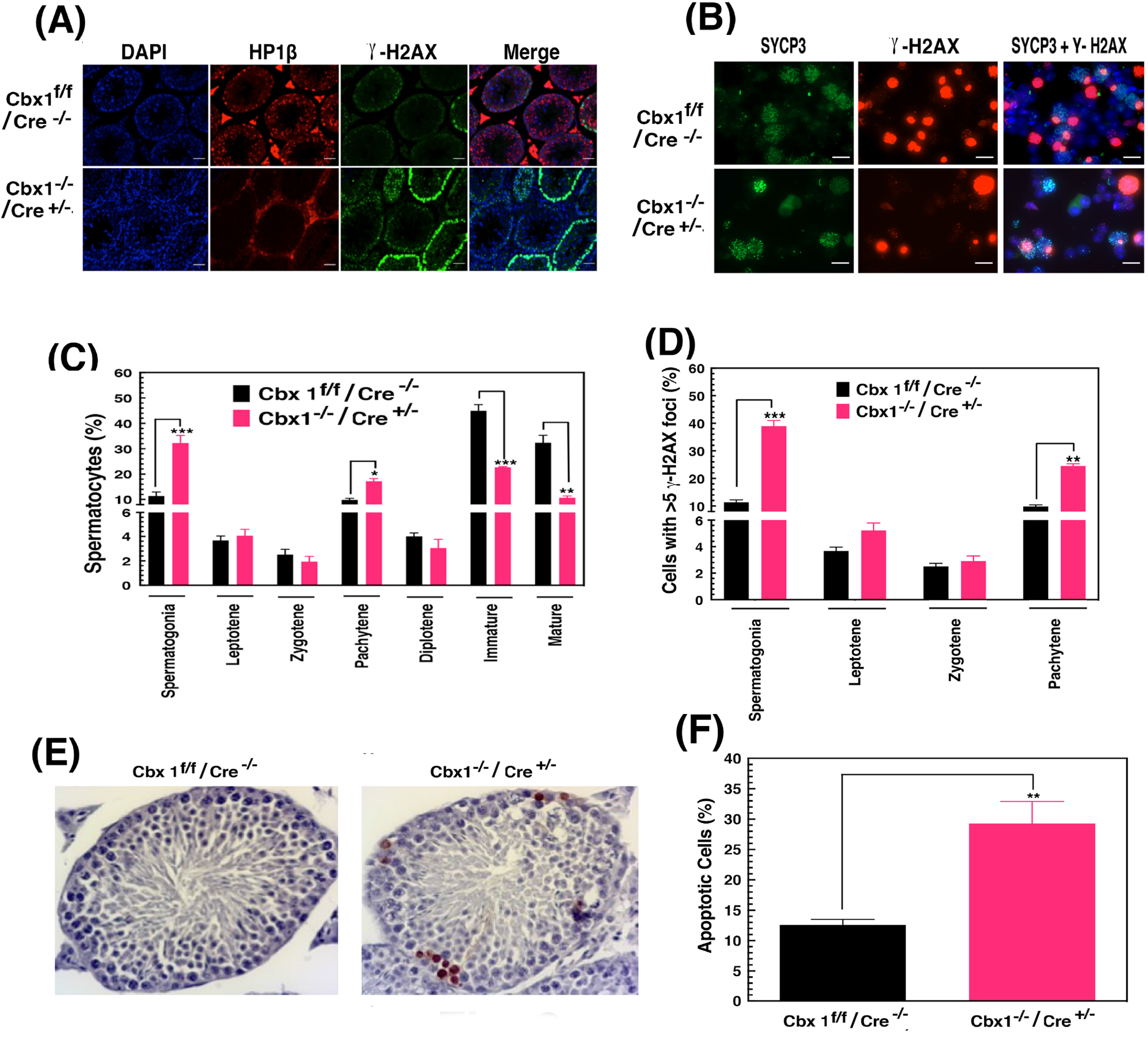
Absence of *Cbx*1 increased DNA damage and apoptosis in testis. **(a)** Levels of HP1β and γ-H2AX in *Cbx1* WT and *Cbx1* testis specific conditional KO mice sections were determined by immunofluorescence staining. **(b)** Staining *Cbx1* WT and *Cbx1* testis specific conditional KO mice testes with anti-SYCP3 antibody observed each sub-stage of meiosis-I.**(c)** Quantification of spermatocytes percentage distribution was analyzed by SYCP3 staining in *Cbx1* WT and *Cbx1* testis specific conditional KO mice. **(d)** Spermatocytes with more than 5 γ-H2AX foci in *Cbx1* WT and *Cbx1* testis specific conditional KO mice were analyzed. **(e)** Tunnel assay conducted in *Cbx1* WT and *Cbx1* testis specific conditional KO mice. **(f)** Quantitification of percentage of apoptotic cells in testes tissue sections. Data analyzed and represented as mean ± SEM. ** P<0.01, Student’s t-test.

### Elevated DNA damage and apoptosis levels in spermatocytes lacking HP1β

HP1β plays an important role in gene repression and DNA repair (Bosch-Presegue et al., 2017; Horikoshi et al., 2019; Kalousi et al., 2015; Kumar et al., 2012; Legartova et al., 2019; Lomberk et al., 2012). The level of endogenous DNA damage in seminiferous tubules of *Cbx1* testis specific conditional KO and *Cbx1* WT testes was measured by γ-H2AX staining, a marker for DNA double strand breaks. In *Cbx1* testis specific conditional KO there was an elevated level of DNA damage in spermatogonia-B cells as well as increased apoptosis in the seminiferous tubules as compared to WT cells (Fig. 2d-f). The increased apoptosis seen in Cbx1 testis specific conditional KO mice was associated with spermatocyte degeneration in the seminiferous tubules (Fig. 1f).

### Role of *Cbx1* in replication fork progression

The *in-vivo* studies suggested that HP1β function could be critical in replicating cells and therefore the DNA replication process was studied in mouse embryonic fibroblasts (MEFs). *Cbx*1^f/f^ MEF cells were prepared and *Cbx*1^−/−^ MEFs subsequently generated by infection with adeno-cre lenti virus (Fig. 3a). Deletion of *Cbx*1 decreased cell proliferation as determined by cell count compared to *Cbx*1^f/f^ MEF cells (Fig. 3b). To determine whether the decreased cell growth and increased DNA damage observed in *Cbx*1^−/−^ MEFs was due to defective DNA replication, we compared replication fork progression in *Cbx*1^−/−^ and *Cbx*1^f/f^ MEFs by fiber analysis and observed that *Cbx*1 deletion increased the number of stalled forks and reduced fork speed (Fig. 3c-f). *Cbx*1^−/−^ MEFs had a reduced nascent fork length, from 12 μm to 7 μm (Fig. 3d), and the fork speed decreased from 1.02 Kb/min to 0.7 Kb/min (Fig. 3e). Loss of *Cbx*1 in MEFs increased the number of endogenous stalled replication forks by 3-fold as determined by fiber assay (Fig. 3f) and basal level DNA damage as evidenced by increased staining for γ-H2AX foci (Fig. 3g).

**Figure 3.**
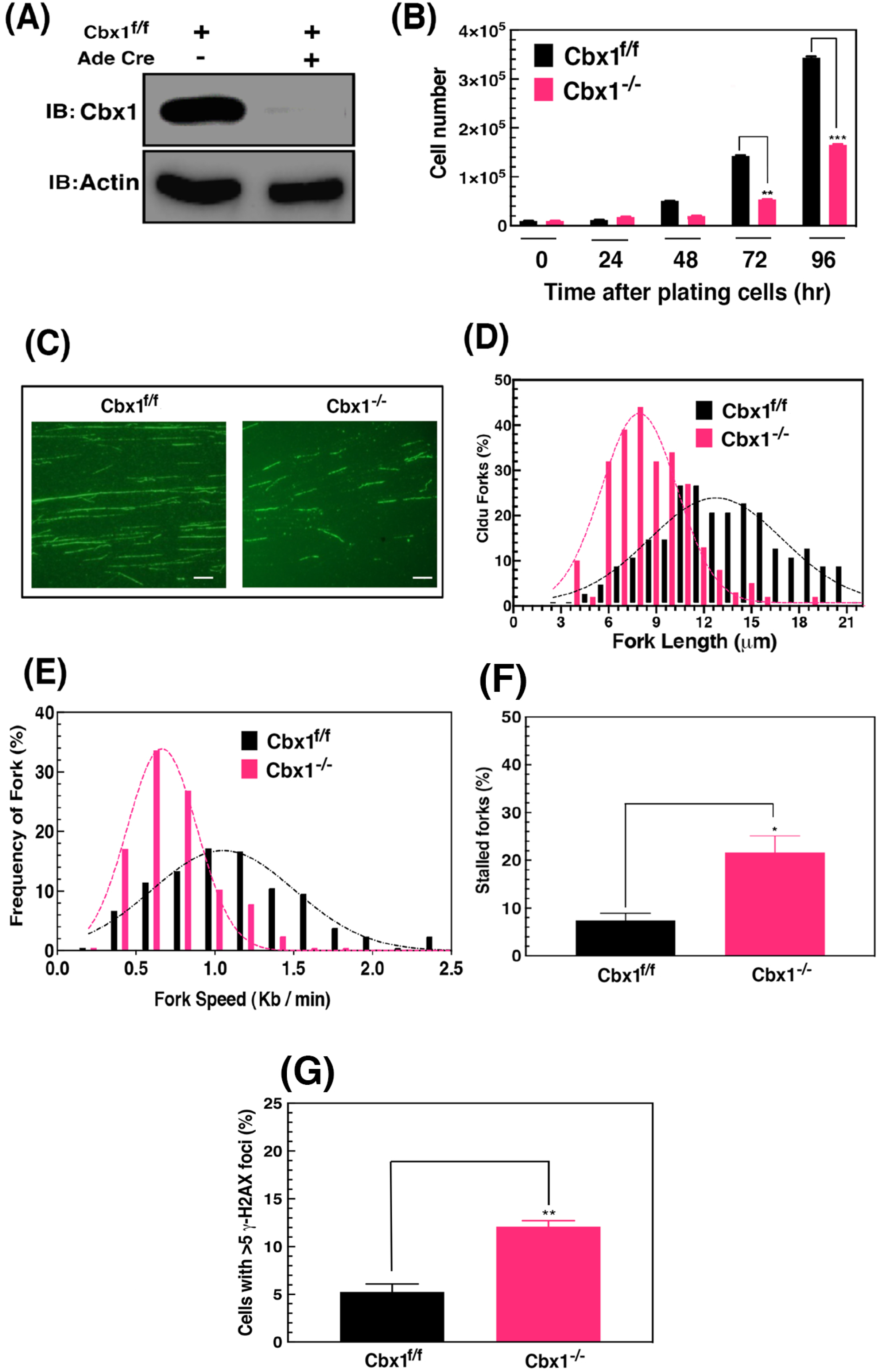
*Cbx*1^−/−^ MEFs exhibit endogenous defect in DNA replication fork progression and stalled fork restart. **(a)** *Cbx*1^−/−^ MEF cell lines confirmed by Western blot analysis. **(b)** Proliferation of *Cbx*1^f/f^,*Cbx*1^−/−^ MEF cells were quantitated by WST assay. **(c-f)**. DNA replication forks progression, fork speed and stalled DNA replication forks in *Cbx*1^−/−^ and *Cbx*1^f/f^ MEFs without hydroxyurea treatment was analyzed. **(g)** *Cbx*1^f/f^ and *Cbx*1^−/−^ MEF cells with more than 5 γ-H2AX foci were analyzed. Average Mean ± SD of 3 independent experiments are shown in. *p < 0.05; **p < 0.01, ***p < 0.001 Student t-test.

To determine whether increased number of stalled forks in HP1β ◻◻◻◻◻◻ed cells could be due to replication fork dynamics, we measured fork progression. Cells were first treated with chlorodeoxyuridine (CldU) to label ongoing DNA replication, washed out and then iododeoxy-uridine (IdU) incorporated to identify replication restarts and new origins (Mattoo et al., 2017). Replication fork dynamics in cells with and without loss of HP1β was measured after treatment with hydroxyurea (HU), which stalls replication by depleting the nucleotide pool. *Cbx*1^−/−^ MEFs or depletion of HP1β with small interfering RNA (siRNA) in HeLa cells have an increased percentage of stalled forks and new origin firings (Fig. 4a-g).

**Figure 4.**
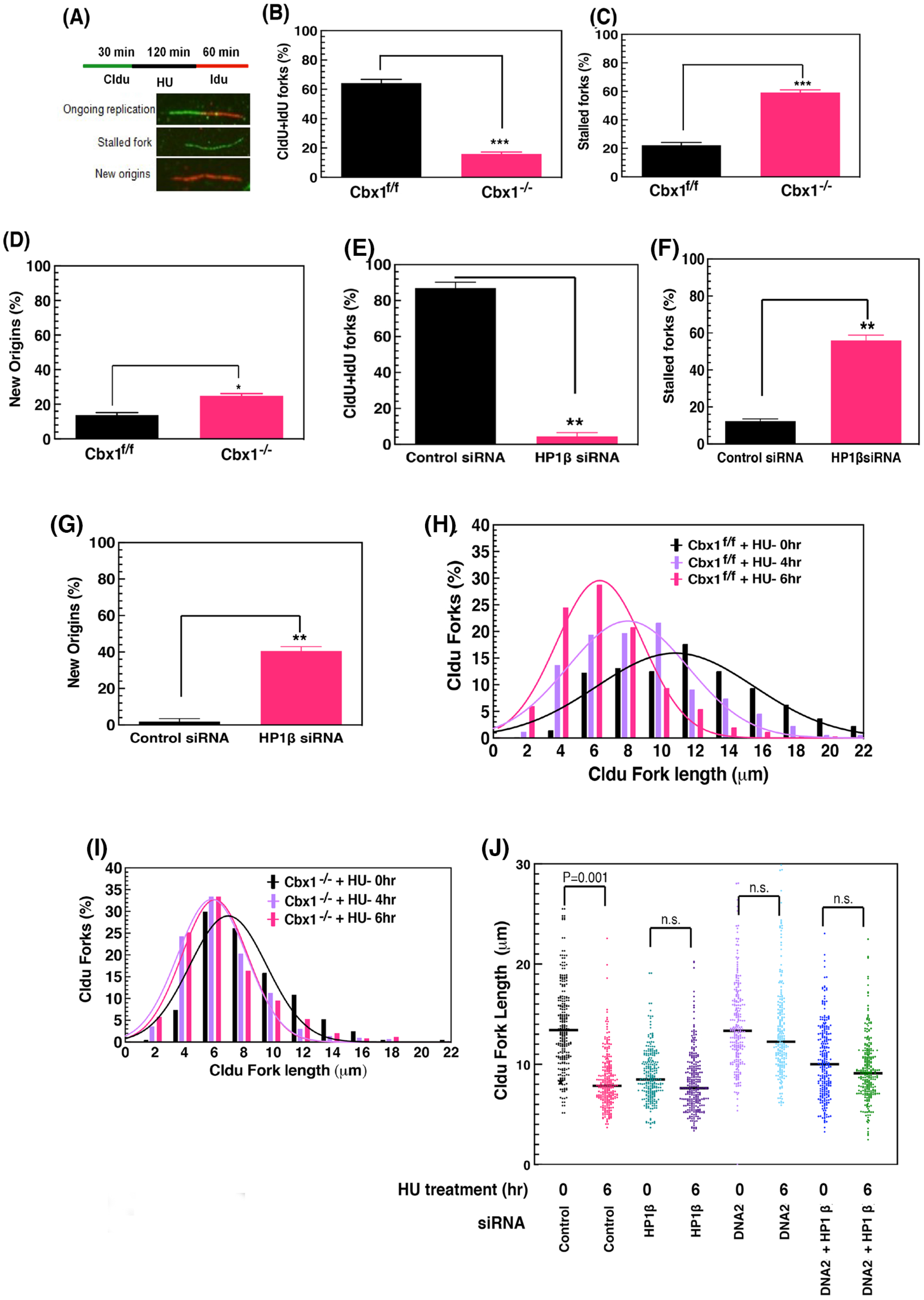
Increased DNA replication fork stalling and new origins in *Cbx*1^−/−^ MEFs and HP1β depleted HeLa cells after hydroxyurea treatment. **(a)** Schematic representation, cells labeled with CIdU, treated with hydroxyurea (HU), IdU for the indicated intervals. **(b,e)** Percentages of recovered stalled DNA replication forks (recovered forks) in *Cbx*1 MEFs and HP1β depleted HeLa cells. **(c,f)** Percentage of stalled forks (CldU nascent DNA) in above cells quantitated. **(d,g)**. New origins (ldU nascent DNA) in in *Cbx*1 MEFs and HP1β depleted cells. Time dependent analysis of nascent replicative DNA fork degradation in **(h,i)** *Cbx*1^f/f^,*Cbx*1^−/−^ MEF cells and in **(j)** HP1β, DNA2, and HP1β/DNA2 depleted HeLa cells. Average Mean ± SD calculated from 3 independent experiments. *p < 0.05; **p < 0.01, ***p < 0.001 Student t-test.

Deletion of *Cbx*1 also decreased restart of fork progression after HU treatment. These un-restarted forks have collapsed, resulting in DNA double strand breaks and collapsed or stalled forks are repaired by HR.

We determined role of HP1β in fork stability in *Cbx*1 MEFs under replicative stress by using DNA fiber assay. For this, cells were labeled with CldU for 30 mins, followed by 2mM HU treatment. DNA fibers were prepared at 0h, 4h and 6h after recovery from HU treatment. Control cells at 0h had an average mean nascent DNA strand fork length of 11.5 μm and the length gradually decreased to 6.2 μm by 6h post HU treatment (Fig. 4h). Cells with *Cbx*1 deletion had a reduced mean average nascent DNA fork length, from 8.3μm to 6.15μm after HU treatment (Fig. 4I). These results suggest HP1β is important for nascent DNA fork progression and stability.

To determine the role of HP1β in fork degradation, we screened for interacting DNA repair proteins. HP1β contains two conserved domains, namely a chromo domain (CD) and a chromo shadow domain (CSD). The CSD recognizes and interacts with proteins containing a PXVXL motif and we data searched for potential HP1β interacting (PxVxL motif containing) proteins. The analysis identified 2916 human PxVxL motif containing proteins, including exonuclease proteins involved in fork degradation (Table. 2).

Among the exonulease proteins, DNA2 is important as it is involved in fork degradation (Pawlowska et al., 2017; Rossi et al., 2018; Thangavel et al., 2015; Zheng et al., 2019). Since either HP1β or DNA2 depletion reduces fork degradation, we determined whether the two proteins function in the same fork degradation pathway. Measurement of nascent DNA fork length in HP1β, DNA2 co-depleted cells did not detect a significant impact on fork degradation after HU treatment as compared to cells with individual HP1β or DNA2 depletion (Fig. 4J). These results suggest HP1β and DNA2 function epistatically in fork degradation (Fig. 4).

### *Cbx*1^−/−^ MEFs are sensitive to drug-induced replicative stress

To evaluate the role of *Cbx*1 in response to drug-induced replication stress, we measured cell survival of *Cbx*1^−/−^ and *Cbx*1^f/f^ MEF cells treated with increasing concentrations of HU or cisplatin. Either deletion of *Cbx*1 in MEFs or depletion of HP1β in HeLa cells increased cell death after HU or cisplatin treatment (Fig. 5a-c). Decreased cell survival correlated with increased chromosome aberrations observed at metaphase, we measured chromosomal aberrations at metaphase after drug treatment, which revealed increased S-phase specific chromosomal aberrations (Fig. 5d-g). These results indicate HP1β is important in repairing DNA damage during S-phase cells (Fig. 5).

**Figure 5.**
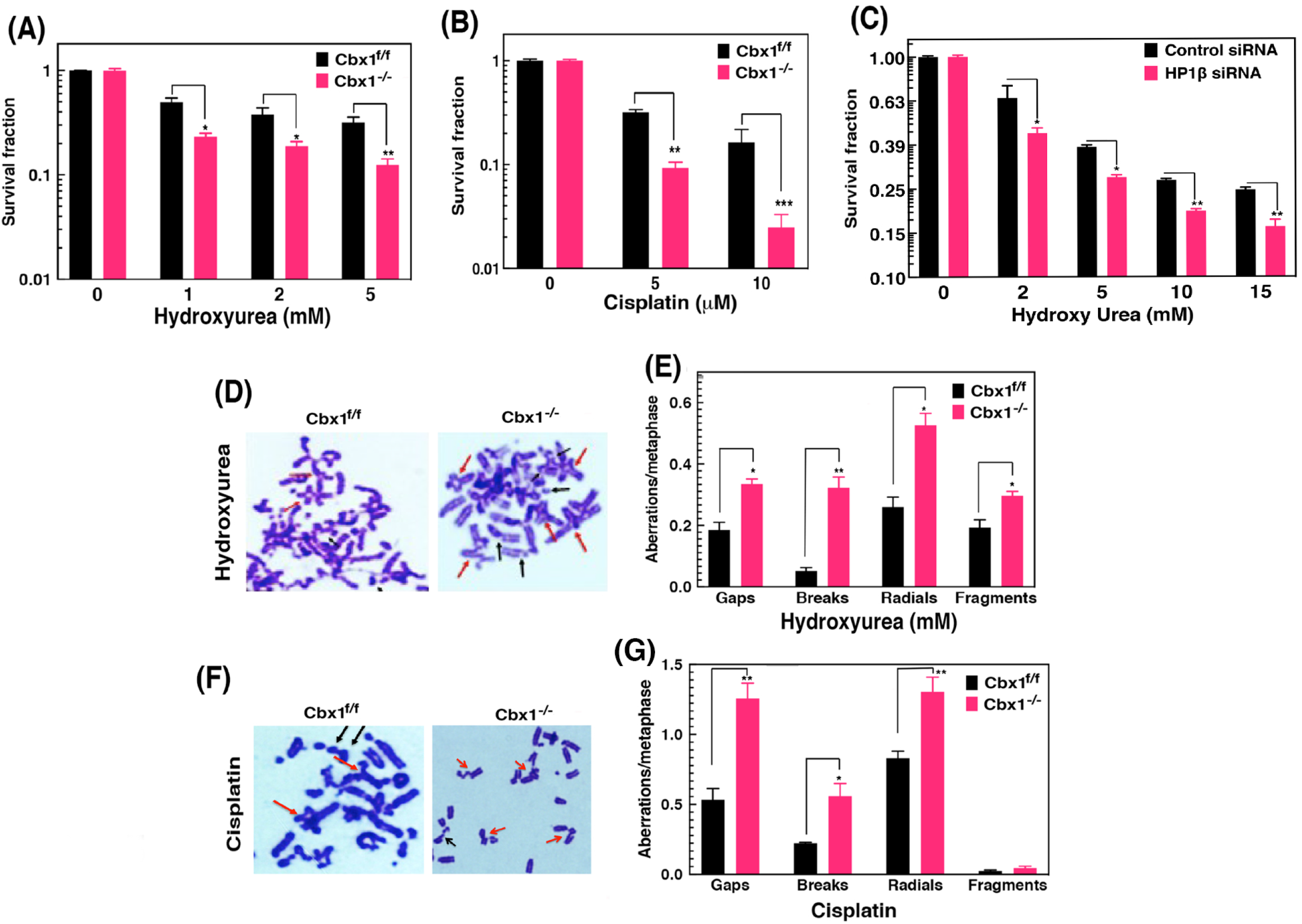
HP1β is essential for cell survival after replication stress. Loss of HP1β has reduced cell survival after exposure to increased concentration of replicative stressors **(a)** Hydroxyurea **(b)** Cisplatin. **(c)** Hydroxy Urea treatment in HP1β depleted HeLa cells (**d, e**) genomic instability with 2mM hydroxyurea (**f, g**) genomic instability with 10μM Cisplatin. Three independent experiments mean ± SD calculated. *p < 0.05; **p < 0.01, ***p < 0.001 Student t-test.

## Discussion

During the germ cell reproductive process, epigenetic chromatin modifications such as methylation, acetylation and ubiquitination play major roles (Gannon et al., 2014). Any alteration in epigenetic modifications could modulate critical gene expression patterns and lead to diseases such as male infertility and failure of embryonic development (Aston et al., 2012; Cho et al., 2003). Histone H3K9me3 and H4K20me3 are associated with heterochromatin formation (Bosch-Presegue et al., 2017). Heterochromatin proteins (HP1) binds to H3K9me3 regions and maintains genomic organization (Santos et al., 2005). The evidence points to distinct functions for HP1β in epigenetic gene regulation of gene expression and epigenetic modifications do impact on DAN damage repair by HR (Bosch-Presegue et al., 2017; Horikoshi et al., 2019; Kalousi et al., 2015; Kumar et al., 2012; Legartova et al., 2019; Lomberk et al., 2012).

To assess the physiological function of HP1β, we generated *Cbx1* testis specific conditional KO mouse because homozygous *Cbx*1^−/−^ neonates die due to severe alveoli damage of lungs and other developmental defects of cerebral cortex (Aucott et al., 2008). HP1β plays role in DNA damage repair during HR and *Cbx1* testis specific conditional KO mouse had defects in spermatogenesis similar to those reported in null mutations for genes involved in DNA repair such as 53BP1, FanCJ, BRCA1, Rnf8, MOF (Guo et al., 2018; Simhadri et al., 2014; Sun et al., 2016; Ward et al., 2003).

Based on the fact that HP1β has a role in DNA repair, specifically during replication stress, the physiological impact of HP1β depletion on germ cells may be related to this mechanism. The manifestation of reduced testicular weight and size in *Cbx1* testis specific conditional KO mouse as well as increased tubular vacuolation, partial degeneration of spermatocytes and decrease sperm production may reflect defective DNA repair in a rapid growing stem cell. The possible reason for sperm reduction in testis with *Cbx*1 deletion is due to higher frequency o γ-H2AX staining levels in spermatogonia B cells and unresolved DNA damaged cells apparently got arrested and subsequently apoptosis, thus resulting in reduction of sperm production and sub-fertility, similar phenotype observed in mice deficient for ATM, PTIP, BRCA1, FANCD2 (Jiang et al., 2018; Schwab et al., 2013; Simhadri et al., 2014)

Our results are consistent with the current literature that HP1β is involved in the DSB repair (Kalousi et al., 2015; Lee et al., 2013; Sharma et al., 2003), and in addition it plays role in the resolution of the stalled replication forks. *Cbx*1 deletion in MEF cells decreased fork speed progression and reduced cellular proliferation compared to wild type MEF cells, thus supporting the argument that like depletion of MOF or FanCD2, HP1β loss also resulted in slow fork progression without any stress (Singh et al., 2018; Zhu et al., 2015). Similar to loss of MOF, FANCD2 and other HR proteins, HP1β depletion increased stalled forks and new origins in HU or Cisplatin treated cells, supporting a role for HP1β in HR repair, which is also critical for germ cell development in order to maintain the fidelity of the DNA. Based on the *in vitro* fiber assay and *in vivo* DNA damage analysis, the potential reason for aberrant spermatogenesis is due to replication stress caused by the absence of HP1β, a novel function for the protein.

## Materials and methods

### Ethics statement

*Cbx1*^tm1a^ KOMP–CSD allele mice were obtained from European consortium. Gt(ROSA)26 Sor tm1(FLP) DYM / RainJ and Tg(Str8-icre)1Reb/J (FLPeR) mice (Farley et al., 2000), mice obtained from Jackson Laboratory. All animal experiments were performed according to guidelines of the Institutional Animal Care Committee of the Houston Methodist Research Institute, USA

### Cell isolation

*Cbx1* WT and *Cbx1* testis specific conditional KO mice were sacrificed by cervical dislocation; testes were excised and washed with PBS. Isolated testis were cut into small pieces and re-suspended in 2ml of TIM buffer containing 40 mg of collagenase. After 1hr, samples were incubated with 1mg/ml trypsin (T8003; Sigma) for 20 mins. DMEM+ 10% FBS media was added to neutralize. These single cell suspensions was washed twice with PBS and re-suspended in 0.1M sucrose solution. Next a 20ul of cellular suspension and 65ul of (1%PFA +0.1% triton x-100) spread on slide. Air-dried slides were stored at −80°C until use (Zelazowski et al., 2017).

Immuno-fluorescence staining was performed by incubating with primary antibodies (γ-H2AX, 1:200, 05–636, Millipore);(SYCP3, 1:200, AB15093, Abcam) for overnight, washed with PBS for times, secondary antibodies (anti-rat Alexa Fluor 488-conjugated [1:100 dilution] and anti-mouse Alexa Fluor 568-conjugated [1:100 dilution] antibodies) for 2h. Each slide was washed with PBS for 3 times. Using an Axio Imager 2.0 microscope captured images and analyzed by Image J software.

### Cell survival assay

*Cbx*1^f/f^ and *Cbx*1^−/−^ MEF cells were seeded onto 60-mm dishes in triplicate. Incubated for 16h, then seeded cells were treated with respective drugs like Cisplatin and Hydroxy Urea. Cisplatin was treated for 1h, where as Hydroxy Urea (HU) for 24h.These treated cells were grown for 13 days and the survived colonies were stained and counted.

### DNA replication restart assay and fork protection assay

DNA fiber assay was done as described previously (Singh et al. 2013; Mattoo et al. 2017). Exponentially growing *Cbx*1^f/f^ and *Cbx*1^−/−^ MEF cells were pulse labeled with 150 mM 5-Chlorodeoxy Uridine (CldU) for 30 mins before stalling with 2mM Hydroxy Urea (HU) for 2h,then washed in PBS for 2 times, incubated in medium containing with 450 mM Iodoxy Uridine (IdU) for 1hr (Chakraborty et al., 2018; Horikoshi et al., 2016).

### Replicative fork progression, stability assays

Exponentially growing cells were incubated in fresh medium containing with 50 mM 5-chlorodeoxyuridine (CldU) for 20 mins, washed in PBS three times, cells were incubated in media containing 450mM IdU for 30 mins. **T**hese labeled cells were treated with 2mM HU. DNA fibers were prepared 0h, 4h, 6h, from the above labeled cells (Schlacher et al., 2012). DNA fiber spreads were fixed, washed in PBS for 3 times, DNA fibers was blocked with 5% BSA for 1h. These fibers were incubated for 1h with anti CldU (1:150) and anti IdU(1:150) antibody, washed in PBS, incubated for 1h in secondary anti-rat Alexa Fluor 488-conjugated (1:100) and anti-mouse Alexa Fluor 568-conjugated (1:100) antibodies solution, each slide washed in PBS. Mounted cell images were acquired by using Axio Imager 2.0, DNA fiber length were analyzed by using Image J software. For each data set, about length of 300 nascent CldU labeled fibers was measured. Results were summarized in bar graphs.

### Hematoxylin and Eosin (H&E) staining, immunohistochemistry and TUNEL staining

Harvested testes were immediately fixed in 4% paraformaldehyde for immunohistochemistry after surgery. For H&E staining, testis tissue sections were deparaffinized, rehydrated, stained with H&E (Kumar et al. 2011; Jiang et al. 2018). TUNEL staining was performed on testis sections according to the instructions provided with the cell death detection kit (11684795910, Roche). To reduce experimental variations, *Cbx1* WT and *Cbx1* testis specific conditional KO mice were processed simultaneously. Images were captured using Axio Imager microscope equipped with a CCD camera and analyzed by Image J software.

### Sperm counting

The epididymis and vasa differentia were removed from *Cbx1* WT and *Cbx1* testis specific conditional KO mice, incised several times. These tissues were incubated in 1 ml PBS at 37°C to release sperms. Number of sperms per testis was counted by using hemocytometer.

### Statistical analysis

All data are presented as mean ± standard deviation (SD). The Mann-Whitney test was applied to compare the number of Υ- H2AX foci and SCP3 between *Cbx1* WT and *Cbx1* testis specific conditional KO mice spermatocytes. The student’s t-test was used for TUNEL and sperm count assays.

